# Residue-Residue Mutual Work Analysis of Retinal-Opsin Interaction in Rhodopsin: Implications for Protein-Ligand Binding

**DOI:** 10.1101/711952

**Authors:** Wenjin Li

## Abstract

Energetic contributions at single-residue level to retinal-opsin interaction in rhodopsin were studied by combining molecular dynamics simulations, transition path sampling, and a newly developed energy decomposition approach. The virtual work at an infinitesimal time interval was decomposed into the work components on one residue due to its interaction with another residue, which were then averaged over the transition path ensemble along a proposed reaction coordinate. Such residue-residue mutual work analysis on 62 residues within the active center of rhodopsin resulted in a very sparse interaction matrix, which is generally not symmetric but anti-symmetric to some extent. 14 residues were identified to be major players in retinal relaxation, which is in excellent agreement with an existing NMR study. Based on the matrix of mutual work, a comprehensive network was constructed to provide detailed insights into the chromophore-protein interaction from a viewpoint of energy flow.

## 1 Introduction

G protein-coupled receptors (GPCRs) are a large superfamily of seven transmembrane helix proteins that transduce extracellular signals to intracellular G proteins.^1,2^ They regulate a variety of cellular signaling pathways and have emerged as a major drug target.^3,4^ The visual pigment rhodopsin is probably the most studied GPCR and is responsible for dim-light vision in vertebrates. The dark state of rhodopsin consists of opsin (the rhodopsin apoprotein) and the 11-*cis* retinal chromophore, which is covalently bound to LYS296 in transmembrane helix 7 (H7) via protonated Schiff base (Figure 1A; residues numbered according to bovine rhodopsin).^5^ Upon absorption of light, the 11-*cis* retinal convert instantly (within 200 fs)^6,7^ to its all-*trans* isomer, which is strongly distorted and stores most of the energy from light.^8^ The distorted all-*trans* retinal then relaxes and induces a chain of conformational changes in rhodopsin involving several spectrally distinct intermediates. Induced structural changes ultimately lead an activated rhodoposin, which triggers the remaining visual cascade.^9–12^

**Figure 1:**
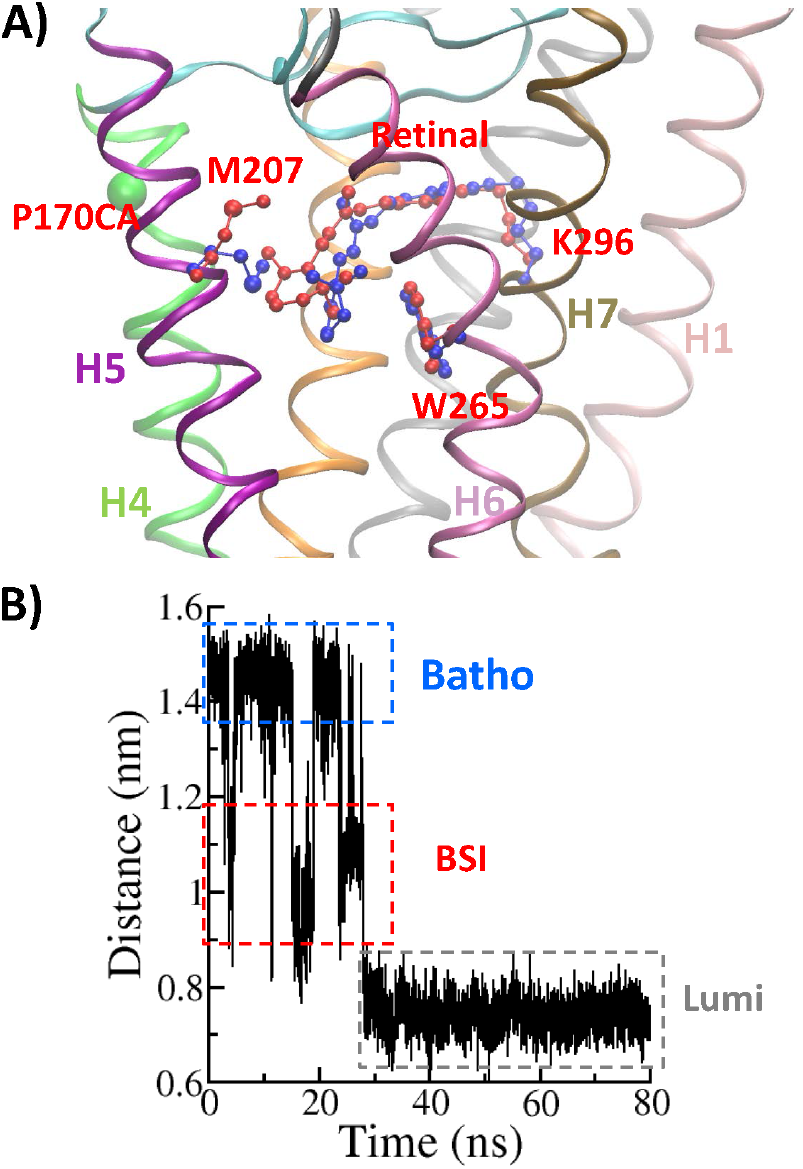
Molecular dynamics from Batho to BSI. A) Comparison between the structures of Batho and BSI. The secondary structure is almost unchanged from Batho to BSI, and thus only the secondary structure of Batho is shown. The residues LYR296, TRP265, and MET207 are shown in ball-stick model, with the ones of Batho highlighted in blue and the ones of BSI in red. The CA atom of the residue PRO170 (P170CA) is shown as a green ball. B) Time evolution of the distance between P170CA and the C4 atom of the retinal. The boxes in blue, red, and grey identify the states of Batho, BSI, and Lumi, respectively.

The photoactivation process of rhodopsin has been intensively studied by both experimental and computational approaches. ^6–8,13–50^ Till now, the structural models of the dark state, intermediates and the activate state are largely known,^24–31^ which formed the basis for numerous computational studies.^32–50^ Computational studies, especially molecular dynamics (MD) simulations in realistic environments and mimicked physiological conditions,^36–50^ provided detailed structural and dynamical information of the conformational changes in the photoactivation process at the atomistic level, however detailed analyses of the change of potential energy terms, especially the energy components involving a single residue, are less commonly touched. The knowledge on the change of energy components at the per-residue or per-coordinate level in the photoactivation process are crucial to answer the questions: which residues are critical for the process and what are their quantitative contributions in the different stages of the transition from one intermediate to the other. Such an information is essential for example in the understanding of protein-ligand interactions and in the development of structural-based pharmacophores with GPCRs as the drug target.

The energetic understanding is limited by the following facts. 1) The current energy decomposition methods, for instance the decomposition of free energy,^51,52^ involves only specific energy terms (or energy components between two atoms or two groups of atoms), they do not distinguish the energy contribution from one group to the other or the other way around; 2) Although methods, e.g., the molecular mechanics-generalized Born surface area (MM/GBSA) approach, can perform per-residue or per-atom decomposition, they usually assume an symmetric contribution of the energy from one group to the other and thus evenly divide the energy term to each group; ^53–55^ 3) The energy decomposition are performed either in a two-end-state way or along an artificial pathway. However, the energy contribution from one group to the other does not equal and simple approximations may result in unreliable results. The dependence of the energy decomposition to the detailed path of integration^56^ suggests that the energy decomposition should be done along the actual path that the reaction follows, that is the reaction coordinate. When the sign is reversed, the change of potential energy can be viewed as the virtual work done to the system. Here, we proposed a method that decomposes the virtual work at an infinitesimal time interval into work components on different residues, which were then averaged over the transition path ensemble (thus termed ensemble averaged work, EAW) along the committor, which was considered to be the “best” reaction coordinate. ^57^ The method is a straightforward extension of an existing method, in which the energy decomposition was performed at the single-coordinate level along the committor in a transition path ensemble. ^58,59^

At the early stage of the photoactivation process in rhodopsin, the first intermediate after light absorption is photorhodopsin, which thermally relaxes within several picoseconds to bathorhodopsin (Batho). On a nanosecond time scale, Batho is in equilibrium with the blue-shifted intermediate (BSI), which decays to form lumirhodopsin (Lumi), an intermediate with a lifetime of microseconds.^13–16^ The dynamic details of the process has been well-studied by previous molecular dynamics simulations.^36,38,49^ In this work, we evaluated the contributions of each residues in terms of EAWs during the transition from Batho to BSI, which is the preliminary step towards a thorough understanding of the whole photo-activation process and also of profound implications for protein-ligand interactions. The EAW on a residue was then further broken down into work components for single residues. Such residue-residue mutual work analysis yielded a matrix of mutual work among almost all residues within the active center of rhodopsin, from which a sparse network of mutual works was constructed. The interaction network revealed novel insights into the chromophore-protein interaction. The transition was demonstrated to be mainly driven by the relaxation of the distorted retinal. TRP265, TYR268 and THR118 are the three prime residues that impede the relaxation. Four residues in the close vicinity of the visual pigment formed a cluster of residues with significant mutual works to each other. In addition, we provided fine details on how the surrounding residues participated in the transition by decomposing residue-wise energy contributions into translational, rotational, and vibrational terms.

## 2 Results and Discussions

### Dynamics from Bathorhodopsin to BSI

The transition from Bathorhodopsin to BSI takes place in a ns timescale and brute-force molecular simulations can simulate such transitions at the atomistic level.^5,36,38,39,49^ Starting with the 3D structural model of Bathorhodopsin (PDB ID: 2G87), which is embedded in an explicit lipid bilayer, we simulated the early relaxation process of the retinal. It was suggested that the structures of Batho, BSI and Lumi can be distinguished by the distance between the residue ALA169 and the *β*-ionone ring.^38^ We thus plotted the distance between the CA atom of the residue PRO170 and the C4 atom of the retinal (see Fig. S7 for names of the atoms in the retinal) as a function of time in Fig 1B, from which three distinct states can be easily identified to be Batho, BSI and Lumi. Several transitions between Batho to BSI were observed within the first 30 ns of the simulation before it relaxed to Lumi. The transition from Batho to BSI completes within a time interval that can be as short as 20 ps, it is thus feasible to harvest by transition path sampling (TPS)^60,61^ the transition path ensembles of such transitions (see Methods for simulation details of TPS). By structural analysis of the transition paths from Batho to BSI, the *β*-ionone ring was observed to move away from TRP265 and displaces MET207 from its original place, and meanwhile rotate about 60 degrees along its long axis (see Fig. 1A and S2). The relaxation dynamics and the structures of BSI and Lumi were consistent with previous computational studies. ^36,38^

### Total EAWs on single residues are comparable to *k*_B_*T*

To identify the key residues that participate in the transition from Batho to BSI, we calculated the committor of each transition path in the transition path ensemble with a fitting procedure, ^62^ and then estimated the overall EAWs of LYR296 and its 61 neighboring residues (for the list of these 62 residues see Table S1; LYR296 or K296^R^ represents the residue of LYS296 covalently bound to the retinal) along the committor, which was quantified as *P*_B_, the probability to relax to Lumi first when starting with a given snapshot (see Methods for more details). As shown in Fig. 2A, the EAWs of each residue along the committor are comparable to thermal fluctuations, with the value close to *k*_B_*T*. Only the EAWs of LYR296 are slightly higher than *k*_B_*T* around *P*_B_ = 1. Thus, it is hard to tell which residue is more importantly involved in the transition. As each residue are interacting with many surrounded residues, lipids and water molecules, such complex environment may cause a dramatic fluctuation of the force on each residues and thus the fluctuation of the EAWs.

**Figure 2:**
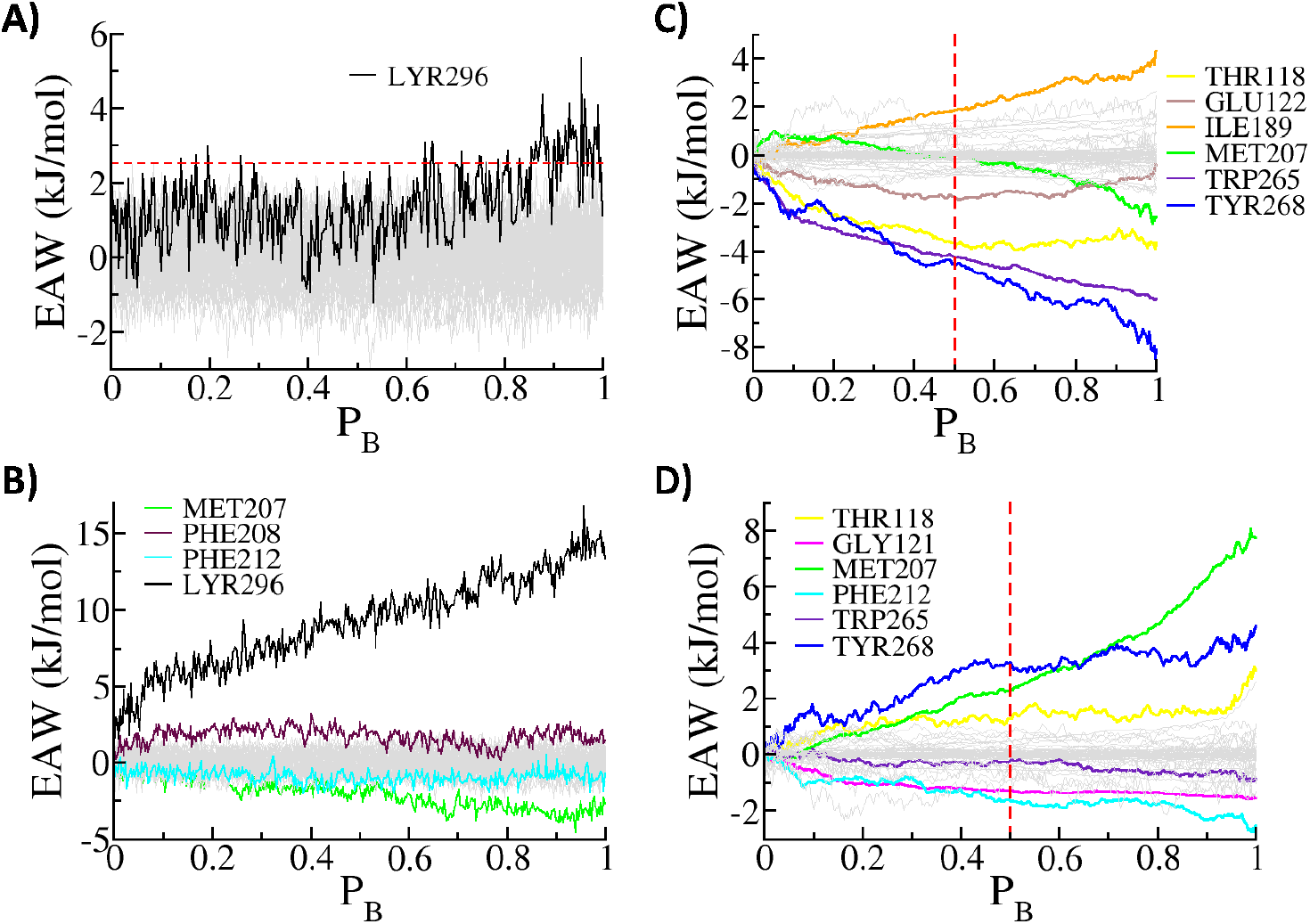
(A) The EAWs of each residue along the committor. Black line: EAW on LYR296; Gray lines: EAWs on all other residues. They are all with the values comparable to *k*_B_*T* or 2.5 kJ/mol (the red dashed line). (B) The EAWs of each residue along the committor due to the change of its internal energy, that is without considering its interaction with the environment. (C) The EAWs of LYR296 along the committor due to its interaction with each neighboring residues. (D) The EAWs of each surrounding residue along the committor due to its interaction with LYR296. In plots B, C, and D, the EAWs of only several residues are highlighted with distinct colors, while the EAWs of all other residues are shown in gray lines. The dashed red lines in plots B and D indicate the values at the transition state where *P*_B_ = 0.5.

### Retinal Relaxes Its Internal Energy

In order to gain clearer results, it is thus helpful to remove the interaction of each residue with its environment, that is to check the EAW of each residue due to the change of its internal energy. We thus excluded the interaction between the residue and their environment and estimated the EAWs of each residue along the committor only due to the interactions within the residue itself (see Fig. 2B). If a residue undergoes significant conformational changes during the transition, the magnitude of EAWs along the committor due to the change of the internal energy will be high. The EAWs of LYR296 is postive and increases to as high as 15 kJ/mol at *P*_B_ = 1, indicating that the distorted retinal is gradually relaxing during the transition. Other residues undergoing obvious conformational changes are MET207, PHE208, and PHE212. While the EAWs of all other coordinates are comparable to 1 kJ/mol, indicating no significant conformational changes.

### Prime residues in retinal relaxation

Comparing the EAWs of LYR296 with and without the environment (see Fig. 2A and 2B), it is obvious that the protein environment have played the role to counterbalance the strong internal energy change of LYR296. To gain insight into the origin of such counterbalancing at a single-residue level, we estimated the EAWs of LYR296 due to its interaction with each surrounding residue (the 61 residues in the list of Table S1 other than LYR296), and the results are shown in Fig. 2C. The counterbalancing EAWs are mainly from residues TYR268, TRP265 and THR118. Around the transition state where *P*_B_ = 0.5, the EAWs from TYR268, TRP265 and THR118 are all about 4 kJ/mol.

When the surrounding residues prevent the relaxation of LYR296, they also receive the work done by LYR296, we thus also estimated the EAWs of each surrounding residue along the committor due to the work done by LYR296 (see Fig. 2D). LYR296 carried out significant amount of work to TYR268 and MET207, but not TRP265, although the work to LYR296 performed by TRP265 was significant (Fig. 2C). Apparently, the works performed to each other for a pair of residues are not necessarily of equal magnitude and opposite sign, although the forces acting to each other are according to Newton’s third law. Note that, MET207 performed negligible work to LYR296 (Fig. 2C), while the work performed by LYR296 to MET207 was *∼*2 kJ/mol at the transition state where *P*_B_ = 0.5. In addition, Residues GLY121, GLU122, ILE189, and PHE212 were involved in the retinal relaxation with smaller but significant contributions.

### Interaction matrix from residue-residue mutual work analysis

The above analysis has revealed the details of EAWs related to LYR296, however the quantitative interaction network between all the neighboring residues can provide further details on how the residues contribute to the relaxation of the retinal. Thus, for all the 62 residues, we estimated the EAWs on one residue due to its interaction with another residue along the committor. Such analysis generated 62*×*62=3844 curves as a function of the committor. For such large number of curves, it is hard to clearly visualize them in the way as shown in e.g., Fig. 2B. As the barrier of the transition is expected to located at the transition state, that is where *P*_B_ = 0.5, we thus took the averaged results in *P*_B_ ∈ [0.45, 0.55] as the EAW on residue *i* due to its interaction with residue *j* as the element [*i, j*] of the interaction matrix around the transition state. The interact matrix is plotted in Fig. 3A. Note that, the results shown in Fig. 2B, 2C, and 2D are the diagonal elements, the 60th column, and the 60th row, respectively. As expected, the interaction matrix is generally not symmetric. Most of the interactions are very small, while the significant interactions occurred among just few residues. Although the works for a pair of residues are not of equal magnitude, it is generally true that they are of opposite sign.

**Figure 3:**
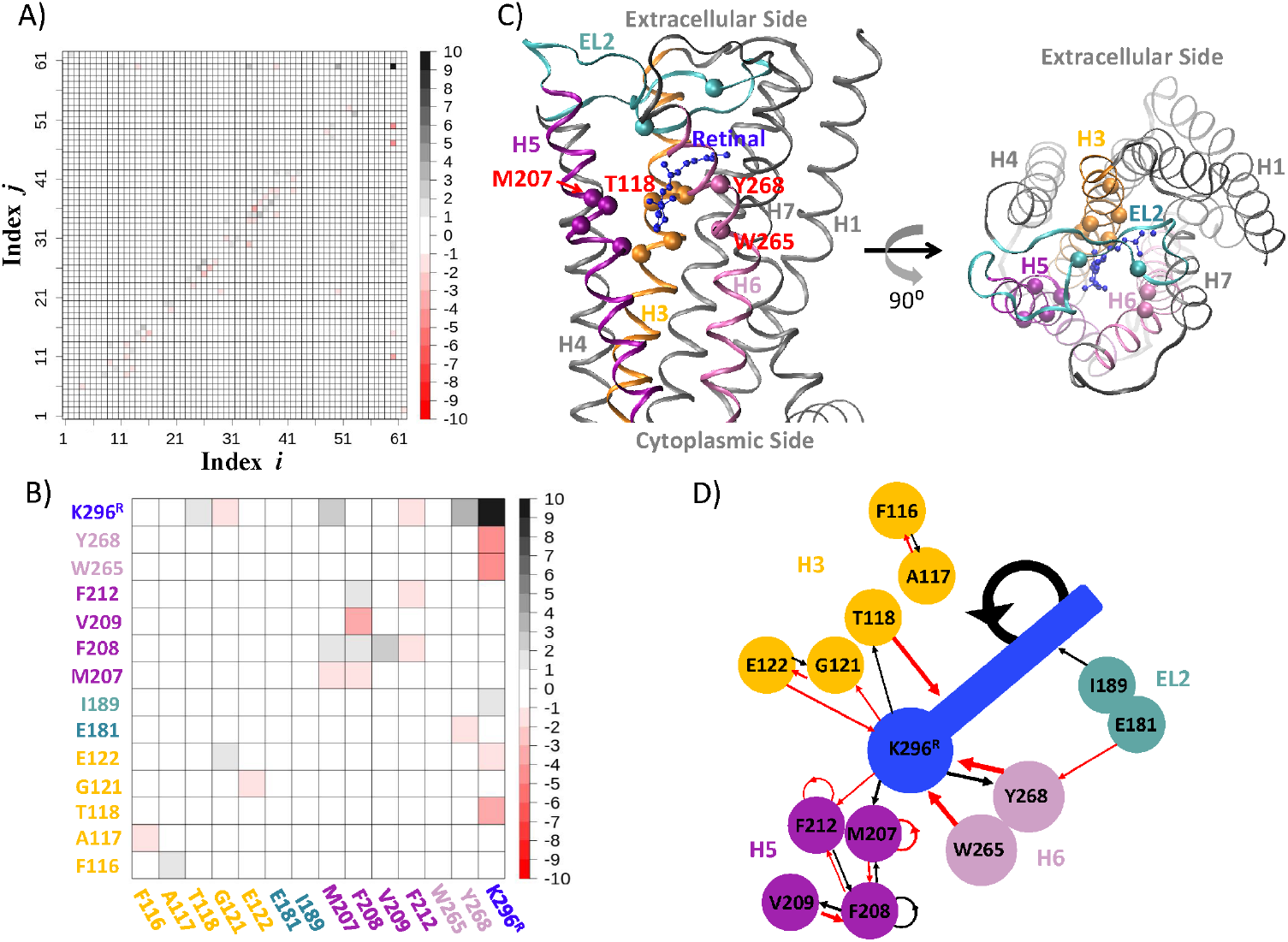
(A) The 62 × 62 matrix of EAWs at the transition state among the 62 residues listed in Table S1. Each element [*i, j*] is the energy contribution from residue *j* to residue *i* (or the work done by residue *j* on residue *i*). (B) The 14 × 14 matrix of EAWs at the transition state among the 14 residues listed in Table S3. For the exact values of the EAWs see Table S5. In plots A and B, color coded according to the scale on the right, which is in unit of kJ/mol. (C) The positions of the selected 14 residues are highlighted in the structure of rhodopsin. The side and top views are shown in left and right, respectively. The CA atoms of all the residues but LYR296 are shown as spheres and colored according to the helix or loop in which they are located. The retinal is shown in ball-stick model in blue. (D) The interaction network of the 14 residues in term of residue-residue mutual works. Residues are in the same colors as the ones in plot C. Positions of residues relative to each other resemble the ones on the right of plot C. The arrow pointed toward a residue indicates the mutual work on the residue. Black and red arrows denote the positive and negative work, respectively. The thickness of an arrow indicates the magnitude of the work. Circular arrow means the mutual work due to the internal energy change of the residue itself.

By assuming that the matrix elements of values larger than 1.5 kJ/mol are significant, we removed the residues whose column and row elements are all less than 1.5 kJ/mol and obtained 26 residues (see Table S2 for the list). To further narrow down the set of main residues involved in the transition, we then summed the elements in row and column of each residue within this 26*×*26 matrix (see Figure S3) and removed 12 residues with negligible work around TS in combination with other criteria (see Figure S4). Thus, the reduced matrix consists of 14 residues (for the list of residues see Table S3) and is shown in Fig. 3B. Note that 10 out of the 14 residues have significant EAWs due to either the change of its internal energy or the interaction with LYR296 (see Fig. 2 and S4). Most of 14 residues were also reported to be important in the activation of rhodopsin in a study combining NMR measurements and MD simulations.^44^

The EAWs along the committor from the interactions among the 62 residues, the EAWs from the interactions among the 26 residues, and the ones among the final 14 residues give similar results (see Fig. S5), indicating that the 14 residues counts for almost all the significant interactions during the retinal relaxation, while all the others play the role of thermal bath.

### Network of mutual works among key residues

The positions of the 14 residues are shown in Fig. 3C, which mainly fall into four groups: TRP265 and TYR268 are on the helix 6 (H6); the MET207, PHE208, VAL209, and HIS211 on H5; the residues PHE116, ALA117, THR118, GLY121, and GLU122 are on H3; and residues GLU181 and THR189 on the extracellular loop 2 (EL2). The 14*×*14 matrix in Fig. 3B is quite sparse as well. There are three main types of mutual works: (1) the work performed to LYR296 by the surrounding residues; (2) the work performed by LYR296 to the surrounding ones; (3) a cluster of mutual works among the residues 207, 208, 209, and 211 on H5. The interaction network of residue-residue mutual work among the 14 residues is then constructed and shown in Fig. 3D. The energy stored in the distorted retinal apparently provides the driving force for the conformational relaxation of the retinal, while TYR268, TRP265, and THR118 are the three major residues to prevent such relaxation. These three residues interact with LYR296 in quite different manners. For example, TRP265 and THR118 interact with LYR296 ‘softly’, in which the work performed to one residue (LYR296) is large while the work performed to the other (TRP265 or THR118) is much smaller and can be even negligible. However, TYR268 interacts strongly with LYR296 in a rather rigid way, in which they perform work to each other with opposite sign and comparable magnitude. The overall change of the interaction energy is thus close to zero and it would be mistakenly regarded as an unimportant residue. These results indicate the necessity to decompose the interactions into residue-residue mutual works.

### Contributions to the translational, rotational, internal movements of the retinal

As TYR268, TRP265, and THR118 are the three main residues that performed significant work to LYR296, we broke down the EAWs on LYR269 from each residues into components resulting from the translational, rotational, internal (or vibrational) movements of the retinal during the transition. The results of such component analysis for TYR268, TRP265, and Thr118 are shown Fig. 4. Although these three residues perform comparable amount of negative work to the retinal, the way that they prevent the relaxation of the retinal are quite different. Both TYR268 and THR118 mainly impede the external movement of LYR296, while TRP265 mostly disfavor the internal organization of LYR296. TYR268 facilitates the translational movement while largely impede the rotational movement of LYR296, while THR118 disfavor the translational and rotational movement equally.

**Figure 4:**
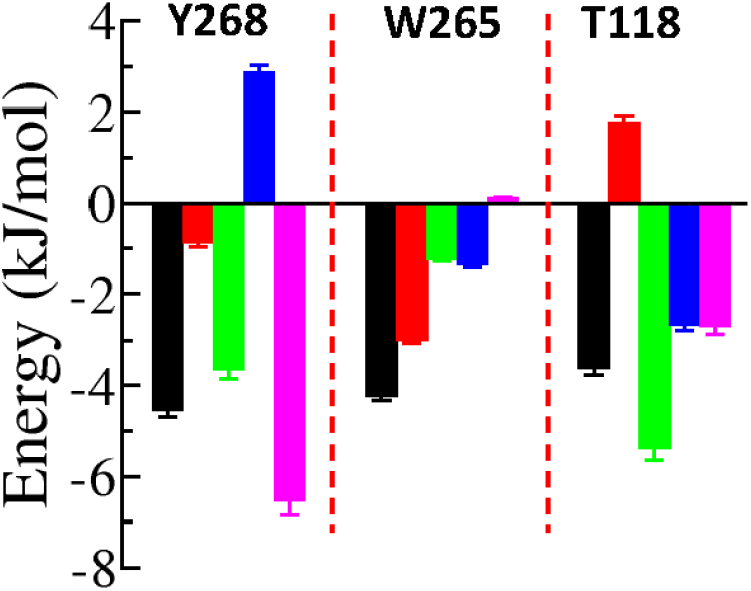
Component analysis of the EAWs on LYR296 from single residues: TYR268 (left), TRP265 (middle), and THR118 (right). The EAW on LYR296 from each residue (black) is decomposed into the internal (red) and external (green) parts. The latter is further broken down into translational (blue) and rotational (purple) components. The error bar is the standard error of the mean.

## 3 Concluding Remarks

MD simulations and transition path sampling were performed to study the dynamics of retinal relaxation in rhodopsin during its transition from Batho to BSI. The observed dynamics was largely consistent with previous computational studies. 14 key residues were determined to be major players in retinal relaxation by a newly proposed energy decomposition method named residue-residue mutual work analysis, in which the work on a residue due to its interaction with another residue was averaged over the transition path ensemble along the committor, a best approximation of the reaction coordinate. Most of these 14 residues were also reported to be fundamental in the activation of rhodopsin from an NMR study.^44^ Thus, residue-residue mutual work analysis could be a reliable approach to identify the dominant contributors in molecular interactions.

Decomposition of the interaction energy between two residues into two mutual works on each other showed its advantage. The net change in interaction energy between two residues can be negligible, while they can perform great amount of mutual works to each other (with opposite sign), as seen in Fig. 3D for LYR296 and TYR268. Such important residues could be missed without the mutual work to each other being estimated.

The resulted matrix of EAWs among the 14 residues was very insightful. It could be general to protein-ligand interactions that the interaction matrix of EAWs is not symmetric but anti-symmetric to some extent. Its sparsity was consist with the general scenario that some key residues dominated the protein-ligand interaction as single-residue mutations in a protein could reduce dramatically the binding affinity of its ligand.

The constructed interaction network provided comprehensive details at single-residue level on how the retinal interacts with the key residues. It would be plausible to propose the following detailed mechanism for the chromophore-protein interaction from Batho to Lumi. All the residues other than the 14 residues were assumed to form the thermal bath of the system. From the viewpoint of energy flow, there were three major ways that the strained energy in retinal were dissipated to bath-mode residues in the thermal bath: (1) A main part of the energy stored in the retinal indirectly passed to bath-mode residues via several key residues, such as TYR268, TRP265, and THR118; (2) A significant part of the energy dissipated indirectly via a group of atoms (the residues MET207, PHE208, VAL209, and HIS211 on H5) that formed a close interaction network; (3) The rest of the energy passed to the thermal bath directly. Here, dissipation means that the EAWs on a residue from another residue is very small and thus can be considered as thermal fluctuations.

EAWs on the retinal from three key residues were further broken down into translational, rotational, and internal components, which provided further insights into the dynamical details of protein-ligand interaction. Residues were observed to act similar overall EAWs on the retinal, but their behaviors were dissimilar on different modes of movements of the retinal. Some of them mainly prevented its overall movement, while others impeded the reorganization of its configurations.

The systematic component analysis at single-residue level could be also carried out to quantify the influences of single-residue mutations on protein-ligand interactions. The resulted interaction matrices and the decomposed work on different modes of movement of the key residues involved in protein-ligand interactions could be beneficial to the construction of structure-based pharmacophore models to enhance virtual screening in computer-aided drug discovery. ^63^ We expected that the applied decomposition scheme can be extended to other biomolecular interactions, for instance, protein-protein interactions.

## 4 Methods

In this section we briefly introduce the MD simulations carried out and the residue-residue mutual work analysis. Further details are provided in Supplemental Methods.

### 4.1 MD Simulations

All simulations were performed with the modified software suite GROMACS-5.1.1^64^ with transition path sampling implemented. The c36 CHARMM force field^65^ was used for the whole system except the retinal, whose force field parameters were adapted from recent literatures.^66,67^ The TIP3P water model was used. The starting structure of Batho was taken from the Protein Data Base (PDB ID 2G87).^27^ Unnecessary heteroatoms were removed and the protonation states of all titratable residues were carefully assigned.^36,68^ The protein was embedded in an explicit lipid bilayer and then hydrated in a box of explicit waters and ions at physiological concentration using the CHARMM-GUI web-server^69^ (see Fig. S6). Proper energy minimization and equilibration were performed before a 80 ns production run to simulate the relaxation of the retinal from Batho to Lumi (see Fig. 1B) in the NPT ensemble. The system was coupled to Berendsen baths to keep bath the temperature and pressure constant.^70^ An integration time step of 2 fs was used with constraints to bonds involving hydrogen atoms using the LINCS algorithm.^71^ The force-switch modifier^72^ and particle mesh Ewald method^73^ were applied for Lennard-Jones interactions and long-range electrostatic interactions, respectively. All molecular structures were visualized with VMD. ^74^

In transition path sampling, the Batho and BSI states were defined by two order parameters: the distance between the CA atom of the residue PRO170 and the C4 atom of the retinal, and the angle formed between the CA atom of the residue GLU122, the C6 and C18 atoms of the retinal. Starting with an initial path taken from the 80 ns production run, 800 independent transition paths were sampled. Transition paths were 20 ps in length. The time step was changed to 0.5 fs, and the method for temperature coupling was changed to the velocity rescaling algorithm.^75^ The committor of a configuration was estimated by shooting 20 trajectories, each trajectory with a maximal length of 100 ps. The fitting procedure recently developed^62^ was applied to evaluating committors of all frames in each path.

### 4.2 Residue-Residue Mutual Work Analysis

The infinitesimal virtual work of a polyatomic system can be decomposed as follows,

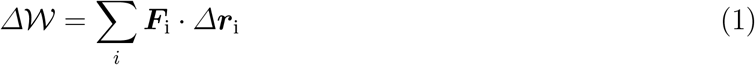

where ***F***_i_ and ***r***_i_ are the force on and the position of atom *i*, respectively. Analogy to an existing approach,^59^ the virtual work on an atom *i* upon an infinitesimal change of the committor *P*_B_ from 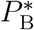 to 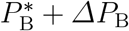 in the transition path ensemble is given by

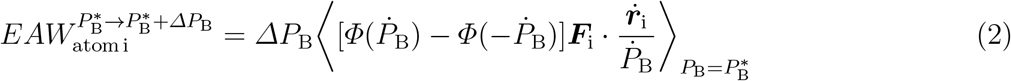

where 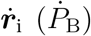 is the time derivative of ***r***_i_ (*P*_B_) and *Φ*(*x*) is a Heaviside step function. 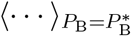 stands for averaging over all the snapshots with 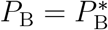 in the transition path ensemble. The term in Eq. 2 is thus named the ensemble averaged work or EAW. The EAW on a residue is then obtained by summing over all its atoms.

In classical MD simulations, the force (***F***_i_) on an atom *i* can be decomposed into pairwise forces between itself and all the other atoms,^76^ that is, 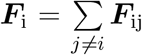 with ***F***_ij_ the pairwise force on atom *i* from atom *j*. Thus, the EAW on a residue can be further broken down into contributions from each residues. The EAW on a residue *k* from a residue *l* is given by,

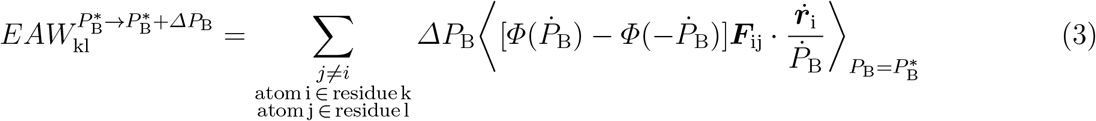

The EAWs along the committor is simply the summation of the EAWs upon each infinitesimal changes of the committor.

It was reported that a complete set of coordinates can be constructed to separate the translational, rotational, and vibrational movements of the system.^77^ Thus, the translational, rotational, and vibrational components of EAWs can be obtained by the summation of EAWs on the corresponding coordinates.

## Supporting information

Details of MD simulations and residue-residue mutual work analysis, Figure S1-S7, and Table S1-S5.

## Acknowledgements

The work was supported by the National Natural Science Foundation of China under Grant No. 31770777 and the Start-up Grant for Peacock Talents, Shenzhen University.

## Supplemental Material

See Supplemental Material for details of MD simulations and residue-residue mutual work analysis, Figure S1-S7, and Table S1-S5.

## References

[1] Fredriksson, R.; Lagerström, M. C.; Lundin, L.-G.; Schiöth, H. B. Molecular pharmacology 2003, 63, 1256–1272.

[2] Rosenbaum, D. M.; Rasmussen, S. G.; Kobilka, B. K. Nature 2009, 459, 356–363.

[3] Pierce, K. L.; Premont, R. T.; Lefkowitz, R. J. Nature Reviews Molecular Cell Biology 2002, 3, 639–650.

[4] Lagerström, M. C.; Schiöth, H. B. Nature reviews Drug discovery 2008, 7, 339–357.

[5] Smith, S. O. Annual review of biophysics 2010, 39, 309–328.

[6] Schoenlein, R.; Peteanu, L.; Mathies, R.; Shank, C. Science 1991, 254, 412–415.

[7] Kukura, P.; McCamant, D. W.; Yoon, S.; Wandschneider, D. B.; Mathies, R. A. Science 2005, 310, 1006–1009.

[8] Kim, J. E.; Tauber, M. J.; Mathies, R. A. Biochemistry 2001, 40, 13774–13778.

[9] Okada, T.; Ernst, O. P.; Palczewski, K.; Hofmann, K. P. Trends in biochemical sciences 2001, 26, 318–324.

[10] Sakmar, T. P.; Menon, S. T.; Marin, E. P.; Awad, E. S. Annual review of biophysics and biomolecular structure 2002, 31, 443–484.

[11] Palczewski, K. Annual review of biochemistry 2006, 75, 743.

[12] Hofmann, K. P.; Scheerer, P.; Hildebrand, P. W.; Choe, H.-W.; Park, J. H.; Heck, M.; Ernst, O. P. Trends in biochemical sciences 2009, 34, 540–552.

[13] Albeck, A.; Friedman, N.; Ottolenghi, M.; Sheves, M.; Einterz, C.; Hug, S.; Lewis, J.; Kliger, D. Biophysical journal 1989, 55, 233.

[14] Hug, S. J.; Lewis, J. W.; Einterz, C. M.; Thorgeirsson, T. E.; Kliger, D. S. Biochemistry 1990, 29, 1475–1485.

[15] Randall, C.; Lewis, J.; Hug, S.; Bjorling, S.; Eisner-Shanas, I.; Ottolenghi, M.; Sheves, M.; Friedman, N.; Kliger, D. Journal of the American Chemical Society 1991, 113, 3473–3485.

[16] Ganter, U. M.; Kashima, T.; Sheves, M.; Siebert, F. Journal of the American Chemical Society 1991, 113, 4087–4092.

[17] Borhan, B.; Souto, M. L.; Imai, H.; Shichida, Y.; Nakanishi, K. Science 2000, 288, 2209–2212.

[18] Ahuja, S.; Hornak, V.; Yan, E. C.; Syrett, N.; Goncalves, J. A.; Hirshfeld, A.; Ziliox, M.; Sakmar, T. P.; Sheves, M.; Reeves, P. J.; Smith, S. O.; Eilers, M. Nature structural & molecular biology 2009, 16, 168–175.

[19] Malmerberg, E.; Bovee-Geurts, P. H.; Katona, G.; Deupi, X.; Arnlund, D.; Wickstrand, C.; Johansson, L. C.; Westenhoff, S.; Nazarenko, E.; Schertler, G. F.; Menzel, A.; de Grip, W. J.; Neutze, R. Science signaling 2015, 8, ra26–ra26.

[20] Elgeti, M.; Rose, A. S.; Bartl, F. J.; Hildebrand, P. W.; Hofmann, K.-P.; Heck, M. Journal of the American Chemical Society 2013, 135, 12305–12312.

[21] Concistre, M.; Gansmüller, A.; McLean, N.; Johannessen, O. G.; Marín Montesinos, I.; Bovee-Geurts, P. H.; Verdegem, P.; Lugtenburg, J.; Brown, R. C.; DeGrip, W. J.; Levitt, M. H. Journal of the American Chemical Society 2008, 130, 10490–10491.

[22] Thomas, Y. G.; Szundi, I.; Lewis, J. W.; Kliger, D. S. Biochemistry 2009, 48, 12283–12289.

[23] Pan, D.; Ganim, Z.; Kim, J. E.; Verhoeven, M. A.; Lugtenburg, J.; Mathies, R. A. Journal of the American Chemical Society 2002, 124, 4857–4864.

[24] Palczewski, K.; Kumasaka, T.; Hori, T.; Behnke, C. A.; Motoshima, H.; Fox, B. A.; Le Trong, I.; Teller, D. C.; Okada, T.; Stenkamp, R. E.; Yamamoto, M.; Miyano, M. science 2000, 289, 739–745.

[25] Okada, T.; Fujiyoshi, Y.; Silow, M.; Navarro, J.; Landau, E. M.; Shichida, Y. Proceedings of the National Academy of Sciences 2002, 99, 5982–5987.

[26] Okada, T.; Sugihara, M.; Bondar, A.-N.; Elstner, M.; Entel, P.; Buss, V. Journal of molecular biology 2004, 342, 571–583.

[27] Nakamichi, H.; Okada, T. Angewandte Chemie International Edition 2006, 45, 4270–4273.

[28] Nakamichi, H.; Okada, T. Proceedings of the National Academy of Sciences 2006, 103, 12729–12734.

[29] Salgado, G. F.; Struts, A. V.; Tanaka, K.; Krane, S.; Nakanishi, K.; Brown, M. F. Journal of the American Chemical Society 2006, 128, 11067–11071.

[30] Choe, H.-W.; Kim, Y. J.; Park, J. H.; Morizumi, T.; Pai, E. F.; Krauß, N.; Hofmann, K. P.; Scheerer, P.; Ernst, O. P. Nature 2011, 471, 651–655.

[31] Scheerer, P.; Park, J. H.; Hildebrand, P. W.; Kim, Y. J.; Krauß, N.; Choe, H.-W.; Hofmann, K. P.; Ernst, O. P. Nature 2008, 455, 497–502.

[32] Gascón, J. A.; Sproviero, E. M.; Batista, V. S. Accounts of chemical research 2006, 39, 184–193.

[33] Campomanes, P.; Neri, M.; Horta, B. A.; Röhrig, U. F.; Vanni, S.; Tavernelli, I.; Rothlisberger, U. Journal of the American Chemical Society 2014, 136, 3842–3851.

[34] Schreiber, M.; Sugihara, M.; Okada, T.; Buss, V. Angewandte Chemie International Edition 2006, 45, 4274–4277.

[35] Khrenova, M.; Bochenkova, A.; Nemukhin, A. Proteins: Structure, Function, and Bioinformatics 2010, 78, 614–622.

[36] Röhrig, U. F.; Guidoni, L.; Rothlisberger, U. Biochemistry 2002, 41, 10799–10809.

[37] Pitman, M. C.; Grossfield, A.; Suits, F.; Feller, S. E. Journal of the American Chemical Society 2005, 127, 4576–4577.

[38] Neri, M.; Vanni, S.; Tavernelli, I.; Rothlisberger, U. Biochemistry 2010, 49, 4827–4832.

[39] Saam, J.; Tajkhorshid, E.; Hayashi, S.; Schulten, K. Biophysical journal 2002, 83, 3097–3112.

[40] Röhrig, U. F.; Guidoni, L.; Laio, A.; Frank, I.; Rothlisberger, U. Journal of the American Chemical Society 2004, 126, 15328–15329.

[41] Röhrig, U. F.; Sebastiani, D. The Journal of Physical Chemistry B 2008, 112, 1267–1274.

[42] Sun, X.; Ågren, H.; Tu, Y. The Journal of Physical Chemistry B 2014, 118, 10863–10873.

[43] Crozier, P. S.; Stevens, M. J.; Forrest, L. R.; Woolf, T. B. Journal of molecular biology 2003, 333, 493–514.

[44] Hornak, V.; Ahuja, S.; Eilers, M.; Goncalves, J. A.; Sheves, M.; Reeves, P. J.; Smith, S. O. Journal of molecular biology 2010, 396, 510–527.

[45] Feng, J.; Brown, M. F.; Mertz, B. Biophysical journal 2015, 108, 2767–2770.

[46] Valsson, O.; Campomanes, P.; Tavernelli, I.; Rothlisberger, U.; Filippi, C. Journal of chemical theory and computation 2013, 9, 2441–2454.

[47] Martínez-Mayorga, K.; Pitman, M. C.; Grossfield, A.; Feller, S. E.; Brown, M. F. Journal of the American Chemical Society 2006, 128, 16502–16503.

[48] Leioatts, N.; Mertz, B.; Martínez-Mayorga, K.; Romo, T. D.; Pitman, M. C.; Feller, S. E.; Grossfield, A.; Brown, M. F. Biochemistry 2014, 53, 376–385.

[49] Lemáıtre, V.; Yeagle, P.; Watts, A. Biochemistry 2005, 44, 12667–12680.

[50] Grossfield, A.; Pitman, M. C.; Feller, S. E.; Soubias, O.; Gawrisch, K. Journal of molecular biology 2008, 381, 478–486.

[51] Boresch, S.; Archontis, G.; Karplus, M. Proteins: Structure, Function, and Bioinformatics 1994, 20, 25–33.

[52] Brady, G. P.; Szabo, A.; Sharp, K. A. J. Mol. Biol. 1996, 263, 123–125.

[53] Gohlke, H.; Kiel, C.; Case, D. A. Journal of molecular biology 2003, 330, 891–913.

[54] Zoete, V.; Meuwly, M.; Karplus, M. Proteins: Structure, Function, and Bioinformatics 2005, 61, 79–93.

[55] Hou, T.; Wang, J.; Li, Y.; Wang, W. Journal of chemical information and modeling 2010, 51, 69–82.

[56] Smith, P. E.; van Gunsteren, W. F. The Journal of Physical Chemistry 1994, 98, 13735–13740.

[57] Li, W.; Ma, A. Molecular simulation 2014, 40, 784–793.

[58] Li, W.; Ma, A. The Journal of chemical physics 2016, 144, 114103.

[59] Li, W.; Ma, A. The Journal of chemical physics 2016, 144, 134104.

[60] Dellago, C.; Bolhuis, P. G.; Csajka, F. S.; Chandler, D. The Journal of chemical physics 1998, 108, 1964.

[61] Bolhuis, P. G.; Chandler, D.; Dellago, C.; Geissler, P. L. Annual review of physical chemistry 2002, 53, 291–318.

[62] Li, W.; Ma, A. The Journal of chemical physics 2015, 143, 11B603 1.

[63] Cele, F. N.; Ramesh, M.; Soliman, M. E. Drug design, development and therapy 2016, 10, 1365.

[64] Hess, B.; Kutzner, C.; Van Der Spoel, D.; Lindahl, E. Journal of chemical theory and computation 2008, 4, 435–447.

[65] Huang, J.; MacKerell Jr, A. D. Journal of computational chemistry 2013, 34, 2135–2145.

[66] Mertz, B.; Lu, M.; Brown, M. F.; Feller, S. E. Biophysical journal 2011, 101, L17–L19.

[67] Zhu, S.; Brown, M. F.; Feller, S. E. Journal of the American Chemical Society 2013, 135, 9391–9398.

[68] Fahmy, K.; Jäger, F.; Beck, M.; Zvyaga, T. A.; Sakmar, T. P.; Siebert, F. Proceedings of the National Academy of Sciences 1993, 90, 10206–10210.

[69] Jo, S.; Kim, T.; Iyer, V. G.; Im, W. Journal of computational chemistry 2008, 29, 1859–1865.

[70] Berendsen, H. J.; Postma, J. v.; van Gunsteren, W. F.; DiNola, A.; Haak, J. The Journal of chemical physics 1984, 81, 3684–3690.

[71] Hess, B.; Bekker, H.; Berendsen, H. J.; Fraaije, J. G. Journal of computational chemistry 1997, 18, 1463–1472.

[72] Steinbach, P. J.; Brooks, B. R. Journal of computational chemistry 1994, 15, 667–683.

[73] Essmann, U.; Perera, L.; Berkowitz, M. L.; Darden, T.; Lee, H.; Pedersen, L. G. The Journal of chemical physics 1995, 103, 8577–8593.

[74] Humphrey, W.; Dalke, A.; Schulten, K. Journal of molecular graphics 1996, 14, 33–38.

[75] Bussi, G.; Donadio, D.; Parrinello, M. The Journal of chemical physics 2007, 126, 014101.

[76] Ishikura, T.; Hatano, T.; Yamato, T. Chemical Physics Letters 2012, 539, 144–150.

[77] Li, W.; Ma, A. The Journal of chemical physics 2015, 143, 12B622_1.

