## Supplementary material for "Residue-Residue Mutual Work Analysis of Retinal-Opsin Interaction in Rhodopsin: Implications for Protein-Ligand Binding": Details of MD simulations and residue-residue mutual work analysis, Figure S1-S7, and Table S1-S5.

Wenjin Li

Institute for Advanced Study, Shenzhen University, Shenzhen, China

#### Supplemental Material

##### Contents:

- Supplemental Methods: MD simulations and residue-residue mutual work analysis
- Supplemental Figures and Tables: Figures S1-S7, Tables S1-S5.

### Supplemental Methods

#### MD Simulations

**Molecular System** A molecular model was constructed based on the high-resolution (2.6 Å) crystal structure of Batho (chain A of PDB ID 2G87).[1] ASP83 and GLU122 were considered to be neutral[2] while all other charged amino acids were set to default protonation states. The spatial positions of Batho with respect to the lipid bilayer were taken from the Orientations of Proteins in Membranes (OPM) database.[3] All histidines were protonated either in the $N_\epsilon$ position (HIS65, HIS152, HIS195, and HIS278) or in the $N_\delta$ position (HIS100 and HIS211)[4]. The Schiff base linkage between the retinal and LYS296 was positively charged. The retinal and all crystallographic water molecules were preserved, other heteroatoms were removed.

The CHARMM-GUI web-server[5] was used to embed the Batho system in an explicit membrane bilayer, and then immerse the resulted protein-lipid system in a box of explicit waters and ions at physiological concentration. Two palmitoyl chains bound to CYS322 and CYS323 were added to Batho before it was inserted in the membrane, which consisted of 128 palmitoyl-oleoyl-phosphatidyl choline (POPC) lipids. The final system was neutral and the size of the periodic box was approximately $71\times71\times92$ Å$^3$ with 8589 water molecules, 20 sodium ions, and 18 chloride ions (see Fig. S6).

**Simulation Details** All simulations were performed with the software suite GROMACS-5.1.1.[6] The c36 CHARMM force field[7] was used for protein and lipids. The force field parameters of retinal were adapted from recent literatures.[8,9] The TIP3P water model was used. The energy of the system was first minimized with very strong position restraints to heavy atoms of the protein (including retinal and two palmitoyl chains bound to rhodopsin). The position of the phosphorus atom and two dihedrals of lipids were also restrained strongly. Then, the system was under a series of equilibration with position and dihedral restraints were gradually released. The early stage of equilibration was performed in the NVE ensemble, then all the subsequent simulations were switched to the NPT ensemble. The system was coupled to a Berendsen thermostat with $\tau_{\mathrm{t}}$=1.0 ps to keep the temperature constant at 303.15 K, and a semiisotropic Berendsen barostat with $\tau_{\mathrm{p}}$=5.0 ps to keep the pressure constant at 1 bar.[10] An integration time step of 2 fs was used with constraints to bonds involving hydrogen atoms using the LINCS algorithm.[11] Lennard-Jones interactions were calculated using a 1.2 nm cutoff with the interaction smoothly switched off starting at 1.0 nm.[12] For electrostatic interactions, the particle mesh Ewald method[13] was applied with a real space cutoff of 1.2 nm. The Verlet cutoff scheme was used for both lennard-Jones and electrostatic interactions. Starting with the equilibrated system, a 80 ns simulation was performed to simulate the relaxation of the retinal from Batho to Lumi (see Fig. 1B in the main text). All molecular structures were visualized with VMD.[14]

**Transition Path Sampling** The Batho and BSI states were defined by two order parameters: the distance ($d$) between the CA atom of the residue PRO170 and the C4 atom of the retinal, and the angle ($\alpha$) formed between the CA atom of the residue GLU122, the C6 and C18 atoms of the retinal. The Batho state was defined as $1.4\,\mathrm{nm} < d < 1.6\,\mathrm{nm}$ and $10° < \alpha < 35°$; The BSI state was defined as $0.95\,\mathrm{nm} < d < 1.2\,\mathrm{nm}$ and $80° < \alpha < 100°$. A transition from Bath to BSI in the 80 ns production run was taken as the initial path and 800 independent transition paths were subsequently sampled with a modified version of GROMACS-5.1.1, in which transition path sampling was implemented. Transition paths were 20 ps in length. The time step was changed to 0.5 fs, and the method for temperature coupling was changed to the velocity rescaling algorithm.[15] Several examples of transition paths were plotted in Fig. S1. The committor of a configuration was estimated by shooting 20 trajectories, each trajectory with a maximal length of 100 ps. The fitting procedure recently developed[16] was applied to evaluating committors of all frames in each path.

#### Residue-Residue Mutual Work Analysis

Emergent potential energy was previously reported to quantify the relevance of a coordinate to the reaction coordinate.[17] For a system of $n$ atoms with $\boldsymbol{q} \equiv (q_1, \cdots, q_{3\mathrm{n}})$ being a complete and independent set of configuration coordinates, an infinitesimal change in the potential energy $\mathcal{V}(\boldsymbol{q})$ of the system can be decomposed as follows,

$$\Delta\mathcal{V}(\boldsymbol{q}) = \sum_i \frac{\partial\mathcal{V}(\boldsymbol{q})}{\partial q_\mathrm{i}}\Delta q_\mathrm{i} \tag{S1}$$

The change of the potential energy on a coordinate $q_\mathrm{i}$ is thus assumed to be $\frac{\partial\mathcal{V}(\boldsymbol{q})}{\partial q_\mathrm{i}}\Delta q_\mathrm{i}$. Then, the emergent potential energy (EPE) on $q_\mathrm{i}$ upon an infinitesimal change of the committor $P_\mathrm{B}$ from $P_\mathrm{B}^*$

to $P_\mathrm{B}^* + \varDelta P_\mathrm{B}$ in the transition path ensemble is given by[17]

$$EPE_{q_\mathrm{i}}^{P_\mathrm{B}^* \to P_\mathrm{B}^* + \varDelta P_\mathrm{B}} = \varDelta P_\mathrm{B} \left\langle [\varPhi(\dot{P}_\mathrm{B}) - \varPhi(-\dot{P}_\mathrm{B})] \frac{\partial \mathcal{V}(\boldsymbol{q})}{\partial q_\mathrm{i}} \frac{\dot{q}_\mathrm{i}}{\dot{P}_\mathrm{B}} \right\rangle_{P_\mathrm{B}=P_\mathrm{B}^*} \tag{S2}$$

where $\dot{q}_\mathrm{i}$ ($\dot{P}_\mathrm{B}$) is the generalized velocity of $q_\mathrm{i}$ ($P_\mathrm{B}$) and $\varPhi(x)$ is a Heaviside step function. $\langle \cdots \rangle_{P_\mathrm{B}=P_\mathrm{B}^*}$ stands for the average over all the snapshots with $P_\mathrm{B} = P_\mathrm{B}^*$ in the transition path ensemble.

Since work equals the negative of a change in potential energy, the ensemble averaged work (EAW) on $q_\mathrm{i}$ when the committor changes is defined as

$$EAW_{q_\mathrm{i}}^{P_\mathrm{B}^* \to P_\mathrm{B}^* + \varDelta P_\mathrm{B}} = -EPE_{q_\mathrm{i}}^{P_\mathrm{B}^* \to P_\mathrm{B}^* + \varDelta P_\mathrm{B}} = \varDelta P_\mathrm{B} \left\langle [\varPhi(\dot{P}_\mathrm{B}) - \varPhi(-\dot{P}_\mathrm{B})] F_{q_\mathrm{i}} \frac{\dot{q}_\mathrm{i}}{\dot{P}_\mathrm{B}} \right\rangle_{P_\mathrm{B}=P_\mathrm{B}^*} \tag{S3}$$

where $F_{q_\mathrm{i}}$ is the generalized force on $q_\mathrm{i}$ and equals $-\frac{\partial \mathcal{V}(\boldsymbol{q})}{\partial q_\mathrm{i}}$. If Cartesian coordinates are used, the EAW on an atom $i$ is given by,

$$EAW_{\mathrm{atom\,i}}^{P_\mathrm{B}^* \to P_\mathrm{B}^* + \varDelta P_\mathrm{B}} = \varDelta P_\mathrm{B} \left\langle [\varPhi(\dot{P}_\mathrm{B}) - \varPhi(-\dot{P}_\mathrm{B})] \boldsymbol{F}_\mathrm{i} \cdot \frac{\dot{\boldsymbol{r}}_\mathrm{i}}{\dot{P}_\mathrm{B}} \right\rangle_{P_\mathrm{B}=P_\mathrm{B}^*} \tag{S4}$$

where $\boldsymbol{F}_\mathrm{i}$ is the force on atom $i$, $\boldsymbol{r}_\mathrm{i}$ and $\dot{\boldsymbol{r}}_\mathrm{i}$ are the position and velocity of atom $i$, respectively. The EAW on a residue $k$ is the summation of EAWs over all the atoms of the residue, that is,

$$EAW_{\mathrm{k}}^{P_\mathrm{B}^* \to P_\mathrm{B}^* + \varDelta P_\mathrm{B}} = \sum_{\mathrm{atom\,i} \in \mathrm{residue\,k}} EAW_{\mathrm{atom\,i}}^{P_\mathrm{B}^* \to P_\mathrm{B}^* + \varDelta P_\mathrm{B}} \tag{S5}$$

In classical MD simulations, the force ($\boldsymbol{F}_\mathrm{i}$) on an atom $i$ can be decomposed into pairwise forces between itself and all the other atoms:[18]

$$\boldsymbol{F}_\mathrm{i} = \sum_{j \neq i} \boldsymbol{F}_\mathrm{ij} \tag{S6}$$

where $\boldsymbol{F}_\mathrm{ij}$ is the pairwise force on atom $i$ from atom $j$. Thus, the EAW on a residue can be further broken down into components from single residues. From Eq. (S4), (S5), and (S6), the EAW on a residue $k$ from a residue $l$ is obtained by,

$$EAW_{\mathrm{kl}}^{P_{\mathrm{B}}^{*}\to P_{\mathrm{B}}^{*}+\Delta P_{\mathrm{B}}} = \sum_{\substack{j\neq i \\ \mathrm{atom\,i}\in\mathrm{residue\,k} \\ \mathrm{atom\,j}\in\mathrm{residue\,l}}} \Delta P_{\mathrm{B}} \left\langle [\Phi(\dot{P}_{\mathrm{B}}) - \Phi(-\dot{P}_{\mathrm{B}})] \boldsymbol{F}_{\mathrm{ij}} \cdot \frac{\dot{\boldsymbol{r}}_{\mathrm{i}}}{\dot{P}_{\mathrm{B}}} \right\rangle_{P_{\mathrm{B}}=P_{\mathrm{B}}^{*}} \tag{S7}$$

The EAWs along the committor is simply the summation of the EAWs upon each infinitesimal changes of the committor.

It was reported that a complete set of coordinates can be constructed to separate the translational, rotational, and internal movements of the system.[19] Thus, the translational, rotational, and internal components of EAWs can be obtained by the summation of EAWs on the corresponding coordinates.

#### Supplemental Figures and Tables

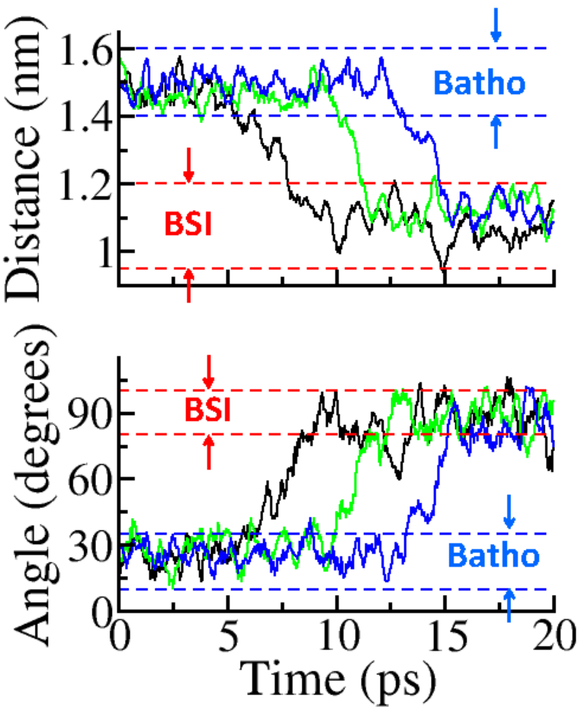


Figure S1: Examples of transition paths are projected onto the two order parameters in transition path sampling. The region of Batho and BSI are indicated by blue and red lines, respectively. Top: the distance between the CA atom of the residue PRO170 and the C4 atom of the retinal; Bottom: the angle formed between the CA atom of the residue GLU122, the C6 and C18 atoms of the retinal.

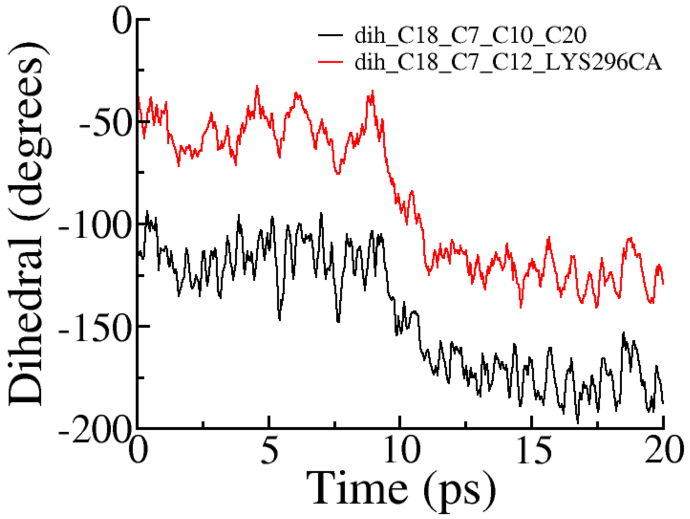


Figure S2: Two dihedrals are used to represent the overall rotation of the retinal along its long axis. The retinal was observed to rotate ∼60 degrees in the transition from Batho to BSI. Black: the dihedral formed between the C18, C7, C10, and C20 of the retinal; Red: the dihedral formed between the C18, C7, and C12 of the retinal, and the CA atom of LYS296.

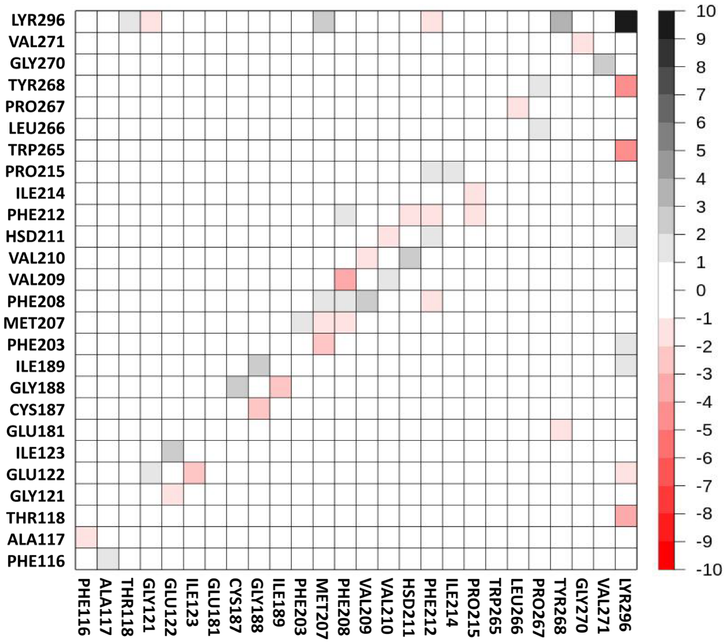


Figure S3: The 26×26 matrix of EAWs at the transition state among the 26 residues listed in Table S2. The values of the elements in the matrix can be found in Table S4. Color coded according to the scale on the right, which is in unit of kJ/mol.

Table S1: The list of 62 residues.

| | | | | | | | |
|---|---|---|---|---|---|---|---|
| 1 | TYR43 | 21 | TYR178 | 41 | ILE214 | 61 | THR297 |
| 2 | MET86 | 22 | GLU181 | 42 | PRO215 | 62 | SER298 |
| 3 | PHE91 | 23 | CYS185 | 43 | LEU216 | | |
| 4 | THR94 | 24 | SER186 | 44 | PHE261 | | |
| 5 | PHE103 | 25 | CYS187 | 45 | LEU262 | | |
| 6 | GLU113 | 26 | GLY188 | 46 | CYS264 | | |
| 7 | GLY114 | 27 | ILE189 | 47 | TRP265 | | |
| 8 | PHE115 | 28 | ASP190 | 48 | LEU266 | | |
| 9 | PHE116 | 29 | TYR191 | 49 | PRO267 | | |
| 10 | ALA117 | 30 | PHE203 | 50 | TYR268 | | |
| 11 | THR118 | 31 | VAL204 | 51 | ALA269 | | |
| 12 | LEU119 | 32 | ILE205 | 52 | GLY270 | | |
| 13 | GLY120 | 33 | TYR206 | 53 | VAL271 | | |
| 14 | GLY121 | 34 | MET207 | 54 | ALA272 | | |
| 15 | GLU122 | 35 | PHE208 | 55 | PHE273 | | |
| 16 | ILE123 | 36 | VAL209 | 56 | ALA292 | | |
| 17 | LEU125 | 37 | VAL210 | 57 | PHE293 | | |
| 18 | ALA166 | 38 | HSD211 | 58 | PHE294 | | |
| 19 | CYS167 | 39 | PHE212 | 59 | ALA295 | | |
| 20 | ALA168 | 40 | ILE213 | 60 | LYR296 | | |

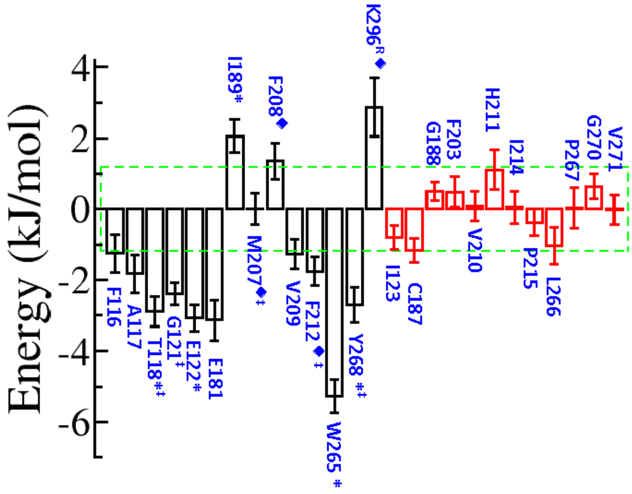

Figure S4: Summing over the elements in the row and column related to a selected residue in the 26×26 matrix (as shown in Fig. S3) yields the total averaged work related to the residue. Each diagonal element is counted once in the summation. A high magnitude of the total averaged work means that the interaction between the residue and its environment underwent a significant change, indicating its important role in the transition from Batho to BSI. Residues whose total average work is of magnitude higher than 1.2 kJ/mol are considered to be important residues. MET207 is considered to be important as the EAW on MET207 from LYR296 is high as shown in Fig. 2D in the main text. Thus, in total 14 residues are selected to be key residues. 11 out of the 14 residues are also observed to be important residues in Fig. 2 in the main text. ♦ indicates the residues with significant changes in their internal energy as shown in Fig. 2B; ∗ indicates the residues that performed significant EAWs on LYR296 as shown in Fig. 2C; ‡ indicates the residues that received significant EAWs from LYR296 as shown in Fig. 2D.

Table S2: The list of 26 residues that are obtained after removing residues with all the row and column elements smaller than 1.5 kJ/mol.

| | | | | | |
|---|---|---|---|---|---|
| 1 | PHE116 | 11 | PHE203 | 21 | LEU266 |
| 2 | ALA117 | 12 | MET207 | 22 | PRO267 |
| 3 | THR118 | 13 | PHE208 | 23 | TYR268 |
| 4 | GLY121 | 14 | VAL209 | 24 | GLY270 |
| 5 | GLU122 | 15 | VAL210 | 25 | VAL271 |
| 6 | ILE123 | 16 | HSD211 | 26 | LYR296 |
| 7 | GLU181 | 17 | PHE212 | | |
| 8 | CYS187 | 18 | ILE214 | | |
| 9 | GLY188 | 19 | PRO215 | | |
| 10 | ILE189 | 20 | TRP265 | | |

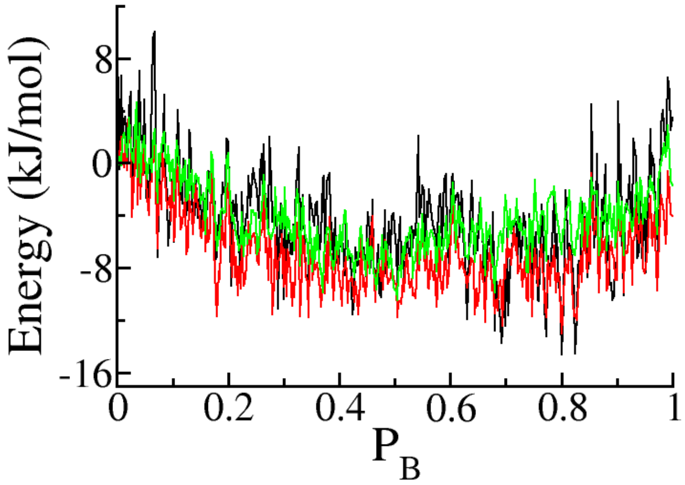


Figure S5: Summed EAWs along the committor resulting from the interactions among 62 residues (black), 26 residues (red), and 14 residues (green). For the lists of residues, see Table S1, S2, and S3, respectively.

Table S3: The list of 14 important residues involved in the transition from Bathorhodopsin to Lumirhodopsin.

| | | | |
|---|---|---|---|
| 1 | PHE116 | 8 | MET207 |
| 2 | ALA117 | 9 | PHE208 |
| 3 | THR118 | 10 | VAL209 |
| 4 | GLY121 | 11 | PHE212 |
| 5 | GLU122 | 12 | TRP265 |
| 6 | GLU181 | 13 | TYR268 |
| 7 | ILE189 | 14 | LYR296 |

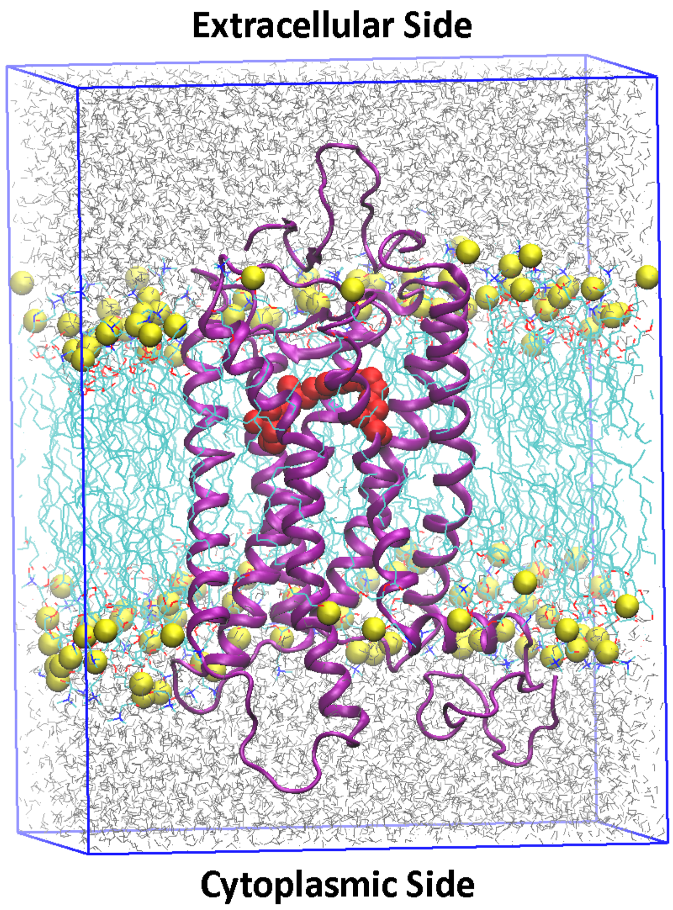


Figure S6: The structural model of Bathorhodopsin (purple) embedded in the POPC lipid bilayer (cyan) and immersed in a box of water (grey) and ions (not shown). The retinal and LYR296 are shown in red. The phosphorus atoms of lipids are shown in yellow spheres. The size of the periodic box is approximately 71×71×92 $\text{Å}^3$ with 8589 water molecules, 128 lipids, 20 sodium ions, and 18 choloride ions.

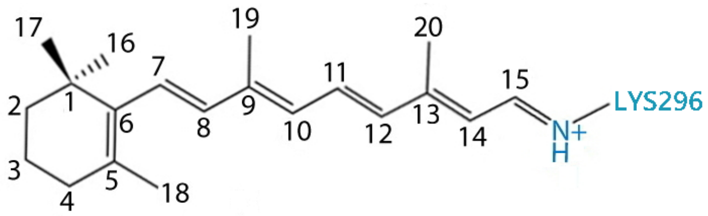


Figure S7: The indices of the carbon atoms in the retinal. Each carbon atom is named as CX, where X is its index.

Table S4: EAWs in the transition state for the 26 residues in Table S2.

| | F116 | A117 | T118 | G121 | E122 | I123 | E181 | C187 | G188 | I189 | F203 | M207 | F208 |
|---|---|---|---|---|---|---|---|---|---|---|---|---|---|
| F116 | -0.4 | -1.9 | -0.1 | 0.0 | -0.2 | -0.1 | 0.1 | 0.0 | 0.1 | 0.0 | 0.0 | 0.0 | 0.0 |
| A117 | 1.2 | -0.4 | 0.1 | 0.0 | 0.1 | -0.1 | 0.2 | 0.1 | 0.0 | 0.0 | 0.0 | 0.0 | 0.0 |
| T118 | 0.2 | 0.2 | -0.7 | -0.3 | -0.5 | 0.0 | 0.3 | 0.1 | -0.1 | 0.1 | -0.1 | -0.1 | 0.0 |
| G121 | -0.1 | 0.0 | 0.5 | -0.5 | 1.6 | -0.2 | 0.1 | -0.1 | 0.0 | 0.0 | 0.0 | 0.0 | 0.0 |
| E122 | 0.1 | -0.1 | 0.3 | -1.8 | -0.9 | 2.3 | 0.0 | 0.0 | 0.0 | 0.1 | -0.1 | -0.1 | 0.0 |
| I123 | 0.0 | 0.2 | 0.0 | 0.1 | -2.2 | -0.8 | 0.0 | 0.0 | 0.0 | 0.0 | 0.0 | 0.0 | 0.0 |
| E181 | -0.1 | 0.0 | -0.1 | 0.0 | 0.0 | 0.0 | -0.8 | -0.3 | 0.1 | 0.1 | 0.0 | 0.0 | 0.0 |
| C187 | 0.0 | 0.0 | 0.0 | 0.0 | 0.0 | 0.0 | -0.1 | -0.5 | 2.6 | 0.1 | 0.0 | 0.0 | 0.0 |
| G188 | -0.1 | 0.0 | 0.0 | 0.0 | 0.0 | 0.0 | -0.8 | -2.5 | 0.5 | 2.4 | 0.0 | 0.0 | 0.0 |
| I189 | 0.0 | 0.0 | 0.0 | 0.0 | -0.1 | 0.0 | 0.2 | -0.1 | -2.5 | 0.0 | -0.5 | -0.4 | 0.1 |
| F203 | 0.0 | 0.0 | 0.0 | 0.0 | 0.1 | 0.0 | 0.0 | 0.0 | 0.0 | 0.7 | 0.2 | 2.0 | 0.1 |
| M207 | 0.0 | 0.0 | -0.2 | 0.0 | -0.4 | 0.0 | -0.1 | 0.0 | 0.0 | 0.8 | -2.7 | -1.9 | 1.7 |
| F208 | 0.0 | 0.0 | -0.1 | 0.0 | -0.1 | 0.0 | -0.1 | 0.0 | 0.0 | 0.0 | -0.3 | -1.4 | 1.9 |
| V209 | 0.0 | 0.0 | 0.0 | 0.0 | 0.0 | 0.0 | 0.0 | 0.0 | 0.0 | 0.0 | -0.1 | 0.3 | 2.5 |
| V210 | 0.0 | 0.0 | 0.0 | 0.0 | 0.0 | 0.0 | 0.0 | 0.0 | 0.0 | 0.0 | 0.1 | 0.1 | 0.1 |
| H211 | 0.0 | 0.0 | 0.0 | -0.1 | -0.4 | 0.0 | 0.0 | 0.0 | 0.0 | 0.0 | 0.2 | -0.2 | 0.3 |
| F212 | 0.0 | 0.0 | 0.1 | -0.1 | -0.2 | -0.1 | 0.0 | 0.0 | 0.0 | 0.0 | 0.1 | 0.4 | -1.2 |
| I214 | 0.0 | 0.0 | 0.0 | 0.0 | -0.1 | 0.0 | 0.0 | 0.0 | 0.0 | 0.0 | 0.1 | 0.0 | 0.0 |
| P215 | 0.0 | 0.0 | 0.0 | 0.0 | 0.1 | 0.0 | 0.0 | 0.0 | 0.0 | 0.0 | 0.0 | -0.1 | 0.1 |
| W265 | 0.0 | 0.2 | 0.1 | -0.5 | -0.2 | -0.1 | 0.0 | 0.0 | 0.0 | 0.0 | 0.0 | 0.0 | 0.0 |
| L266 | 0.0 | 0.0 | 0.0 | 0.0 | 0.1 | 0.0 | 0.0 | 0.0 | 0.0 | 0.0 | 0.0 | 0.0 | 0.1 |
| P267 | 0.0 | 0.0 | 0.0 | 0.0 | 0.0 | 0.0 | 0.1 | 0.0 | 0.0 | 0.0 | 0.0 | 0.0 | 0.0 |
| Y268 | 0.1 | -0.1 | -0.1 | 0.1 | 0.0 | 0.0 | -1.5 | 0.0 | 0.1 | -0.5 | -0.1 | -0.2 | -0.1 |
| G270 | 0.0 | 0.0 | 0.0 | 0.0 | 0.0 | 0.0 | 0.0 | 0.0 | 0.0 | 0.0 | 0.0 | 0.0 | 0.0 |
| V271 | 0.0 | 0.0 | 0.0 | 0.0 | 0.0 | 0.0 | -0.1 | 0.0 | 0.0 | -0.1 | 0.0 | 0.0 | 0.0 |
| K296$^{R}$ | -0.1 | -0.5 | -3.7 | 0.0 | -1.8 | -0.1 | -0.6 | 0.0 | 0.9 | 1.8 | 1.0 | 0.0 | 0.0 |

| | V209 | V210 | H211 | F212 | I214 | P215 | W265 | L266 | P267 | Y268 | G270 | V271 | K296$^{R}$ |
|---|---|---|---|---|---|---|---|---|---|---|---|---|---|
| F116 | 0.0 | 0.0 | 0.0 | 0.0 | 0.0 | 0.0 | 0.0 | 0.0 | 0.0 | 0.0 | 0.0 | 0.0 | 0.0 |
| A117 | 0.0 | 0.0 | 0.1 | 0.0 | 0.0 | 0.0 | -0.1 | 0.0 | 0.0 | 0.1 | 0.0 | 0.0 | -0.8 |
| T118 | 0.0 | 0.0 | 0.0 | -0.1 | 0.0 | 0.0 | -0.1 | 0.0 | 0.0 | 0.1 | 0.0 | 0.0 | 1.4 |
| G121 | 0.0 | 0.0 | 0.0 | 0.0 | 0.1 | 0.0 | 0.3 | 0.0 | 0.0 | -0.1 | 0.0 | 0.0 | -1.3 |
| E122 | 0.0 | 0.2 | -0.1 | 0.1 | 0.1 | -0.2 | 0.2 | 0.0 | 0.0 | 0.0 | 0.0 | 0.0 | 0.7 |
| I123 | 0.0 | 0.0 | -0.1 | 0.0 | 0.0 | 0.0 | 0.0 | 0.0 | 0.0 | 0.0 | 0.0 | 0.0 | 0.1 |
| E181 | 0.0 | 0.0 | 0.0 | 0.0 | 0.0 | 0.0 | 0.0 | -0.1 | 0.0 | 0.1 | 0.0 | 0.0 | 0.4 |
| C187 | 0.0 | 0.0 | 0.0 | 0.0 | 0.0 | 0.0 | 0.0 | 0.0 | 0.0 | -0.1 | 0.0 | 0.0 | -0.4 |
| G188 | 0.0 | 0.0 | 0.0 | 0.0 | 0.0 | 0.0 | 0.0 | 0.0 | 0.0 | 0.0 | 0.0 | 0.0 | -0.1 |
| I189 | 0.0 | 0.0 | -0.1 | 0.0 | 0.0 | 0.0 | -0.1 | 0.0 | 0.0 | -0.2 | 0.0 | 0.1 | 0.3 |
| F203 | 0.0 | -0.1 | 0.0 | -0.1 | -0.1 | 0.0 | 0.0 | 0.0 | 0.0 | 0.1 | 0.0 | 0.0 | 0.0 |
| M207 | -0.2 | -0.1 | -0.1 | 0.1 | -0.1 | 0.2 | 0.1 | 0.1 | 0.0 | 0.1 | 0.0 | 0.1 | 2.4 |
| F208 | -3.0 | -0.1 | -0.1 | 1.5 | 0.1 | -0.1 | -0.1 | -0.1 | 0.0 | 0.0 | 0.0 | 0.0 | -0.3 |
| V209 | 0.1 | -1.8 | 0.2 | -0.1 | -0.2 | -0.1 | 0.0 | 0.0 | 0.0 | 0.0 | 0.0 | 0.0 | 0.0 |
| V210 | 1.2 | 0.3 | -1.6 | 0.1 | 0.1 | -0.1 | 0.0 | 0.0 | 0.0 | 0.0 | 0.0 | 0.0 | 0.0 |
| H211 | 0.0 | 2.2 | -0.5 | -1.4 | 0.0 | 0.0 | 0.0 | -0.1 | 0.0 | 0.0 | 0.0 | 0.0 | 0.1 |
| F212 | -0.4 | -0.3 | 1.0 | -1.1 | 0.2 | 1.2 | 0.4 | 0.3 | 0.0 | -0.1 | 0.0 | 0.0 | -1.6 |
| I214 | 0.3 | -0.1 | 0.0 | 0.0 | -0.2 | 1.4 | 0.0 | 0.0 | 0.0 | 0.0 | 0.0 | 0.0 | -0.1 |
| P215 | 0.0 | 0.0 | 0.3 | -1.1 | -1.6 | -0.4 | 0.0 | 0.0 | 0.0 | 0.0 | 0.0 | 0.0 | 0.0 |
| W265 | 0.0 | 0.0 | 0.0 | -0.2 | 0.0 | 0.0 | -0.2 | -0.6 | -0.1 | 0.6 | 0.0 | 0.1 | -0.3 |
| L266 | 0.0 | 0.0 | 0.1 | -0.1 | 0.0 | 0.0 | -0.2 | 0.2 | -1.5 | 0.2 | -0.3 | 0.0 | -0.1 |
| P267 | 0.0 | 0.0 | 0.0 | 0.0 | 0.0 | 0.0 | 0.0 | 1.2 | -0.3 | 1.1 | 0.0 | -0.2 | 0.1 |
| Y268 | 0.0 | 0.0 | 0.0 | 0.0 | 0.0 | 0.0 | -0.4 | -0.1 | -0.5 | 0.0 | -0.1 | 0.5 | 3.1 |
| G270 | 0.0 | 0.0 | 0.0 | 0.0 | 0.0 | 0.0 | 0.0 | 0.0 | 0.2 | 0.1 | 0.5 | -1.5 | 0.0 |
| V271 | 0.0 | 0.0 | 0.0 | 0.0 | 0.0 | 0.0 | 0.0 | -0.1 | -0.1 | -0.2 | 2.0 | -0.7 | 0.0 |
| K296$^{R}$ | 0.0 | -0.1 | 1.3 | 0.7 | 0.0 | 0.0 | -4.2 | -0.2 | 0.0 | -4.6 | -0.1 | 0.1 | 9.3 |

Table S5: EAWs in the transition state for the 14 residues in Table S3. Elements with EAWs larger than 1 kJ/mol are highlighted in bold.

| | F116 | A117 | T118 | G121 | E122 | E181 | I189 | M207 | F208 | V209 | F212 | W265 | Y268 | $K296^R$ |
|---|---|---|---|---|---|---|---|---|---|---|---|---|---|---|
| $K296^R$ | 0.0 | -0.8 | **1.4** | **-1.3** | 0.7 | 0.4 | 0.3 | **2.4** | -0.3 | 0.0 | -1.6 | -0.3 | **3.1** | **9.3** |
| Y268 | 0.0 | 0.1 | 0.1 | -0.1 | 0.0 | 0.1 | -0.2 | 0.1 | 0.0 | 0.0 | -0.1 | 0.6 | 0.0 | **-4.6** |
| W265 | 0.0 | -0.1 | -0.1 | 0.3 | 0.2 | 0.0 | -0.1 | 0.1 | -0.1 | 0.0 | 0.4 | -0.2 | -0.4 | **-4.2** |
| F212 | 0.0 | 0.0 | -0.1 | 0.0 | 0.1 | 0.0 | 0.0 | 0.1 | **1.5** | -0.1 | **-1.1** | -0.2 | 0.0 | 0.7 |
| V209 | 0.0 | 0.0 | 0.0 | 0.0 | 0.0 | 0.0 | 0.0 | -0.2 | **-3.0** | 0.1 | -0.4 | 0.0 | 0.0 | 0.0 |
| F208 | 0.0 | 0.0 | 0.0 | 0.0 | 0.0 | 0.0 | 0.1 | **1.7** | **1.9** | **2.5** | **-1.2** | 0.0 | -0.1 | 0.0 |
| M207 | 0.0 | 0.0 | -0.1 | 0.0 | -0.1 | 0.0 | -0.4 | **-1.9** | **-1.4** | 0.3 | 0.4 | 0.0 | -0.2 | 0.0 |
| I189 | 0.0 | 0.0 | 0.1 | 0.0 | 0.1 | 0.1 | 0.0 | 0.8 | 0.0 | 0.0 | 0.0 | 0.0 | -0.5 | **1.8** |
| E181 | 0.1 | 0.2 | 0.3 | 0.1 | 0.0 | -0.8 | 0.2 | -0.1 | -0.1 | 0.0 | 0.0 | 0.0 | **-1.5** | -0.6 |
| E122 | -0.2 | 0.1 | -0.5 | **1.6** | -0.9 | 0.0 | -0.1 | -0.4 | -0.1 | 0.0 | -0.2 | -0.2 | 0.0 | -1.8 |
| G121 | 0.0 | 0.0 | -0.3 | -0.5 | **-1.8** | 0.0 | 0.0 | 0.0 | 0.0 | 0.0 | -0.1 | -0.5 | 0.1 | 0.0 |
| T118 | -0.1 | 0.1 | -0.7 | 0.5 | 0.3 | -0.1 | 0.0 | -0.2 | -0.1 | 0.0 | 0.1 | 0.1 | -0.1 | **-3.7** |
| A117 | **-1.9** | -0.4 | 0.2 | 0.0 | -0.1 | 0.0 | 0.0 | 0.0 | 0.0 | 0.0 | 0.0 | 0.2 | -0.1 | -0.5 |
| F116 | -0.4 | **1.2** | 0.2 | -0.1 | 0.1 | -0.1 | 0.0 | 0.0 | 0.0 | 0.0 | 0.0 | 0.0 | 0.1 | -0.1 |

#### References


[1] Nakamichi, H.; Okada, T. *Angewandte Chemie International Edition* **2006**, *45*, 4270–4273.

[2] Fahmy, K.; Jäger, F.; Beck, M.; Zvyaga, T. A.; Sakmar, T. P.; Siebert, F. *Proceedings of the National Academy of Sciences* **1993**, *90*, 10206–10210.

[3] Lomize, M. A.; Pogozheva, I. D.; Joo, H.; Mosberg, H. I.; Lomize, A. L. *Nucleic acids research* **2011**, *40*, D370–D376.

[4] Röhrig, U. F.; Guidoni, L.; Rothlisberger, U. *Biochemistry* **2002**, *41*, 10799–10809.

[5] Jo, S.; Kim, T.; Iyer, V. G.; Im, W. *Journal of computational chemistry* **2008**, *29*, 1859–1865.

[6] Hess, B.; Kutzner, C.; Van Der Spoel, D.; Lindahl, E. *Journal of chemical theory and computation* **2008**, *4*, 435–447.

[7] Huang, J.; MacKerell Jr, A. D. *Journal of computational chemistry* **2013**, *34*, 2135–2145.

[8] Mertz, B.; Lu, M.; Brown, M. F.; Feller, S. E. *Biophysical journal* **2011**, *101*, L17–L19.

[9] Zhu, S.; Brown, M. F.; Feller, S. E. *Journal of the American Chemical Society* **2013**, *135*, 9391–9398.

[10] Berendsen, H. J.; Postma, J. v.; van Gunsteren, W. F.; DiNola, A.; Haak, J. *The Journal of chemical physics* **1984**, *81*, 3684–3690.

[11] Hess, B.; Bekker, H.; Berendsen, H. J.; Fraaije, J. G. *Journal of computational chemistry* **1997**, *18*, 1463–1472.

[12] Steinbach, P. J.; Brooks, B. R. *Journal of computational chemistry* **1994**, *15*, 667–683.

[13] Essmann, U.; Perera, L.; Berkowitz, M. L.; Darden, T.; Lee, H.; Pedersen, L. G. *The Journal of chemical physics* **1995**, *103*, 8577–8593.

[14] Humphrey, W.; Dalke, A.; Schulten, K. *Journal of molecular graphics* **1996**, *14*, 33–38.

[15] Bussi, G.; Donadio, D.; Parrinello, M. *The Journal of chemical physics* **2007**, *126*, 014101.

[16] Li, W.; Ma, A. *The Journal of chemical physics* **2015**, *143*, 11B603_1.

[17] Li, W.; Ma, A. *The Journal of chemical physics* **2016**, *144*, 134104.

[18] Ishikura, T.; Hatano, T.; Yamato, T. *Chemical Physics Letters* **2012**, *539*, 144–150.

[19] Li, W.; Ma, A. *The Journal of chemical physics* **2015**, *143*, 12B622_1.
